# The H3K27 acetyltransferase p300 is dispensable for thermogenic adipose tissue formation and function

**DOI:** 10.1101/2023.08.23.554499

**Authors:** Daniel Gamu, Makenna S. Cameron, William T. Gibson

## Abstract

Brown adipose tissue (BAT) is specialized for thermogenesis because it contains uncoupling protein (UCP)-1. BAT is also an endocrine organ, producing many signalling molecules important for regulating the metabolism of peripheral organs. Mounting evidence suggest that histone modifying enzymes are integral for the development, tissue maintenance, and postnatal functioning of brown and beige adipocytes. p300 and its functional homologue CREB-binding protein (CBP) are histone acetyltransferases that form the transcriptionally activating histone 3 acetyl-lysine 27 (H3K27ac) mark. Using *Ucp1-*Cre, we examined the requirement of p300 activity specifically within thermogenic adipocytes. We hypothesized that loss of p300 activity would impair gene programming integral for BAT development/function, rendering knockouts susceptible to metabolic dysfunction and unable to form beige adipocytes. Despite successful knockdown, brown fat was completely unaffected by p300 deletion. As such, knockout mice showed a comparable metabolic profile to littermate controls in response to diet-induced obesity. Furthermore, *de novo* beige adipogenesis within subcutaneous fat by a β_3_ adrenergic agonist remained intact in knockout mice. Although p300 and CBP have non-overlapping roles in other tissues, our results indicate p300 HAT activity is dispensable within thermogenic fats, likely due to functional compensation by CBP.

## Introduction

Brown adipose tissue (BAT) is specialized for thermogenesis because it contains uncoupling protein (UCP)-1 within its mitochondrial inner membrane. When activated by the sympathetic nervous system, UCP-1 uncouples ATP production from the mitochondrial electrochemical H^+^ gradient, creating heat instead of ATP.^1^ As such, BAT activity is an integral component mammalian adaptive non-shivering thermogenesis, helping to defend body temperature in times of thermal stress. In addition to discrete depots of classical brown fat, rodents have beige (or ‘brite’) adipocytes interspersed within subcutaneous white fat.^2–5^ While similar to brown adipocytes in morphology and positive expression of UCP-1, beige adipocytes arise from a completely distinct cellular lineage.^6^ *De novo* beige adipogenesis can be induced physiologically or pharmacologically.^6^ For example, cold exposure is well known to increase the number beige adipocytes found within white fat of mice (i.e. “browning” of white fat).^6^ Similarly, pharmacological agonism of the β_3_-adrenergic receptor in rodents, a major regulatory node controlling the functional capacity of thermogenic fats,^1^ also enhances white adipose tissue browning.^7–9^

Numerous rodent studies have shown that altering brown/beige fat thermogenesis impacts energy balance and susceptibility to obesity. As such, there has been considerable historical interest in harnessing BAT activity as a means to reduce body weight and improve systemic metabolism. While accepted that humans possess active brown fat past infancy,^10–12^ it is not abundantly present in healthy adults, limiting its contribution to whole-body energy expenditure.^13^ However, BAT activity is correlated with lower cardiometabolic risk factors.^14–16^ It should be noted that brown fat is more than a thermogenic organ. Just like white fat, BAT secretes numerous peptides and metabolites with paracrine and endocrine function termed BATokines, which have various regulatory roles in brown fat and other peripheral tissues.^15^ For example, BAT derived interleukin-6 reduces diet-induced insulin resistance by stimulating glucose uptake into BAT, white fat and the heart.^17^ Furthermore, BAT-derived phospholipid transfer protein^18^ and the lipokine 12,13-diHOME^19,20^ enhance peripheral cholesterol and lipid metabolism, respectively. Such BAT-derived signals may help to mitigate various dysfunctional metabolic features accompanying obesity and type 2 diabetes, independent of thermogenesis. Despite the expanding role brown fat plays in metabolism, our understanding of pathways governing its development, adaptability, lifespan, and various functions are incompletely understood. It has become increasingly clear over the past decade that mechanisms regulating chromatin structure are vital for controlling plasticity of thermogenic adipocytes.

Within chromatin, DNA is wrapped around histones (H), which are important for nuclear DNA packaging and gene expression. Post-translational modifications (PTMs) of histones tighten or loosen chromatin’s conformation, thereby controlling the transcriptional machinery’s access to important regulatory sites of cellular identity. One critical histone residue is H3 lysine 27 (i.e. H3K27), which is either acetylated (H3K27ac) or mono/di/tri-methylated (H3K27me1-3), which are associated with transcriptional activation or repression, respectively.^21^ Inhibiting various histone deacetylases (HDACs), the enzymes that remove acetyl moieties, improves thermogenic functioning of brown/beige fat, mitigating metabolic dysfunction associated with obesity.^22–28^ The metabolic benefits associated with HDAC loss-of-function are presumed to result from a greater occupancy of the transcriptionally activating H3K27ac mark on critical loci controlling BAT functionality. Although studies of HDACs suggest H3K27ac is a critical molecular switch controlling BAT identity and function, they have failed to consider the histone acetyltransferases (HATs) that actively lay down the H3K27ac mark.

Acetylation of H3K27 depends on the activity of CREB-binding protein (CBP) and its functional homologue p300.^29,30^ Owing to the large number of signaling proteins with which they interact,^31^ these ubiquitously expressed HATs fulfill broad-ranging cellular functions. While numerous classes of histone acetyltransferases exist,^32^ p300/CBP are solely responsible for catalyzing H3K27ac formation.^33–35^ Evidence for a role of p300/CBP in thermogenic adipocytes is currently indirect. For example, adrenergic stimulation of BAT with isoproterenol increases binding p300/CBP and concomitant H3K27ac deposition at thermogenic genes, including *Ucp1* and *Pgc1a*.^26,36^ Similarly, HAT binding and H3K27ac enrichment at the *Prdm16* locus, a protein critical for maintaining brown fat identity,^37^ increases as brown adipocytes mature *in vitro*.^38^ Together, these studies suggest that transcriptional activation of gene programs integral for BAT development and thermogenesis require adequate H3K27ac levels; however, it is yet unclear whether this relationship is obligatory, and if so, to what extent. Here, we sought to determine the *in vivo* requirement of p300 specifically in the maintenance and remodeling of thermogenic adipose tissues. We hypothesized that lack of p300 activity would prevent transcriptional activation of thermogenic gene programming, thereby impairing development and adaptability of brown and beige fats.

## Materials and Methods

### Animals

Mice carrying *Cre-recombinase* driven by the *Ucp1* promoter (i.e. *Ucp1-Cre*) were purchased from Jackson Laboratories (Stock No: 024670; B6.FVB-Tg(Ucp1-cre)1Evdr/J), while those with *loxP* sites flanking exon 9 of *ep300* were developed as described previously.^31^ Breeding pairs consisted of mice that were homozygous for the floxed *ep300* gene (i.e. *ep300*^f*l/fl*^); male hemizygous carriers of *Ucp1*-*Cre* were bred with *Cre*-negative females, generating floxed controls (*fl/fl*) and BAT-specific p300 knockout mice (p300^BAT-/-^). Additionally, all mice were homozygous for the dual-fluorescence *Cre*-reporter allele ROSA^mT/mG^ (Jackson Laboratory; strain #: 007576); thus, cell membrane-localized tdTomato (mT) is replaced by membrane-localized green fluorescent protein (mG) in *Cre*-*recombinase* expressing cells. All animals were housed at room temperature (∼22°C), provided water and standard rodent chow *ad libitum* under a 12-hr light/12-hr dark cycle at BC Children’s Hospital Research Institute animal facility. Experiments were conducted with 10-12 week old male and female mice, and were approved by the University of British Columbia Animal Care Committee in accordance with the Canadian Council on Animal Care guidelines.

### Metabolic Phenotyping

Whole-body energy expenditure, O_2_-consumption, CO_2_-production, respiratory exchange ratio (VCO_2_/VO_2_), spontaneous cage activity, food and water consumption were measured at room temperature in the PhenoMaster home-cage metabolic platform (TSE Systems). Data acquisition was recorded over 96-hrs, with the first 24-hr discarded to remove the effects of acute handling stress on mice. Following metabolic cage experiments, acute cold tolerance was examined by lowering housing temperature to 4°C for 6 hrs. Immediately prior to and following the cold challenge, intrarectal temperature was measured with a RET-3 thermocouple probe (Physitemp) and digital thermometer (ThermoWorks). Mice were then housed back at room temperature for one week, after which they were anesthetized with isoflurane and euthanized by cervical dislocation. Adipose tissues were then dissected and cleared of extraneous tissues, snap frozen in liquid nitrogen and stored at -80°C until analysis (described below).

### Diet-Induced Obesity

Beginning at 10-12 weeks of age, group housed mice were provided *ad libitum* access to water and a “Westernized” high-fat diet (TD.88137, Envigo; 42% kcal fat, 48.5% kcal carbohydrate, 15.2% kcal protein) for 16 weeks as we have previously done.^39,40^ Fresh food was presented every other day, with animal body-mass recorded weekly. Body composition was measured in non-anesthetized mice by quantitative magnetic resonance (EchoMRI-100H, Echo Medical Systems) prior to, and after 8 and 16 weeks of high-fat feeding. At these same timepoints, whole-body glucose tolerance was analyzed following a 5 hr fast; mice were given an intraperitoneal injection of glucose (2g/kg body mass), with blood glucose (OneTouch Ultra; Johnson and Johnson) sampled via tail prick at 0, 15, 30, 60, and 90 min post-injection.

### Remodelling of Thermogenic Adipose Tissue by a Selective ß_3_-Receptor Agonist

To induce beige adipogenesis (i.e. “browning” of white fat), mice were given the highly-selective *ß*_3_-adrenergic receptor agonist CL-316,243 (CL; Sigma; 1 mg/kg body mass) dissolved in saline. Mice were weighed and given an intraperitoneal injection of CL or a weight adjusted volume of vehicle control daily for 7 consecutive days. 24 hours following the final injection, mice were euthanized, and adipose tissues collected as described above.

### qPCR and Western Blotting

One portion of snap-frozen adipose tissue was homogenized in TRIzol using a bead mill (Bullet Blender, Next Advance) and 0.5 mm zirconium oxide beads. RNA was then column purified (RNeasy Mini Kit, Qiagen) following on-column DNase digestion (Qiagen, Cat. No 79254) according to manufacturer instructions. 1 µg of total RNA was reverse-transcribed (SuperScript™ III: Invitrogen, Cat. No 18080-051) using oligo(dT)_20_ primers, after which cDNA was stored at -20°C. On the day of use, cDNA was diluted 1:10 in nuclease-free H_2_O, with adipose transcripts amplified with GoTaq® qPCR master mix (Promega, Cat. No A6001) using a ViiA 7 PCR system (Applied Biosystems) set to Standard cycling conditions. Gene expression data was normalized to *Actb* and calculated using the 2-^ΔΔCT^ method. Primer pairs are listed in Supplemental **Table S1**.

A second portion of snap-frozen adipose tissue was homogenized as above in RIPA buffer supplemented with Halt™ protease/phosphatase inhibitor cocktail (Prod. No 1861284, ThermoScientific), including the histone deacetylase inhibitor sodium butyrate (10 mM final concentration). Protein concentration was determined by the BCA assay (ThermoScientific) using bovine serum albumin as standards. Following electrophoresis, samples were wet-transferred onto PVDF membranes (pore size: 0.2 µm) and probed overnight at 4°C for UCP-1 (ab10983, Abcam), mitochondrial respiratory chain subunits (total OXPHOS; ab110413, Abcam), and for the following using primary anti-bodies from Cell Signalling Technologies: green fluorescent protein (GFP; 4B10), acetylated H3K27 (Lys27), H3 (D1H2), and ß-Tubulin (D3U1W). Following incubation in appropriate secondary anti-bodies (CST), immunoblots were visualized on film using enhanced chemiluminescent substrates.

### Statistics

Mean differences between two groups were analysed by a Student’s T-Test (two-tailed, independent samples), whereas all other data were analyzed by two-way ANOVA or two-way repeated measures ANOVA, followed by a Holm-Sidak post-hoc comparisons where appropriate (GraphPad Prism 10.0.0). Statistical significance was considered at *P<*0.05. Values presented are mean ± S.E.M.

## Results

### Metabolic Characterization of p300^BAT-/-^ Mice

Successful *Cre*-mediated recombination in brown fat was noted by a marked induction of GFP expression and concomitant knockdown of *ep300* exon 9 gene expression in p300^BAT-/-^ mice (**Fig. 1 A/B**). Despite p300 depletion from BAT, whole-body energy expenditure and substrate oxidation preference (RER) were comparable at room temperature between control and knockout mice (**Fig. S1A**). Furthermore, feeding/drinking behaviour and spontaneous cage locomotion did not differ between *fl/fl* and p300^BAT-/-^ littermates (**Fig. S1B-D**). We next assessed thermogenic competency by measuring intrarectal temperature following an acute exposure to 4°C, which knockout mice were able to withstand without developing relative hypothermia after 6 hrs of exposure (**Fig. 1C**). Interestingly, female p300^BAT-/-^ mice were better able to maintain body temperature relative to *fl/fl* controls after 6 hrs of thermic challenge. Following our whole-body metabolic characterization, adipose tissues were excised for examination of molecular markers of thermogenesis. Unlike hypothesized, loss of p300 did not impair brown fat formation, as noted by comparable iBAT depot wet weights of adult p300^BAT-/-^ animals (**Fig. 1D**). Additionally, WAT depot masses were unaffected by p300 knockout, further suggesting energy balance was not perturbed in these mice. Surprisingly, we found no impairment in the expression of thermogenic and adipogenic maker genes despite loss of a major transcriptional co-activator (**Fig. 1E**). Correspondingly, iBAT UCP-1 protein content and the transcriptionally activating H3K27ac mark were expressed to a comparable extent between *fl*/*fl* and p300^BAT-/-^ littermates (**Fig. 1F**).

**Figure 1.**
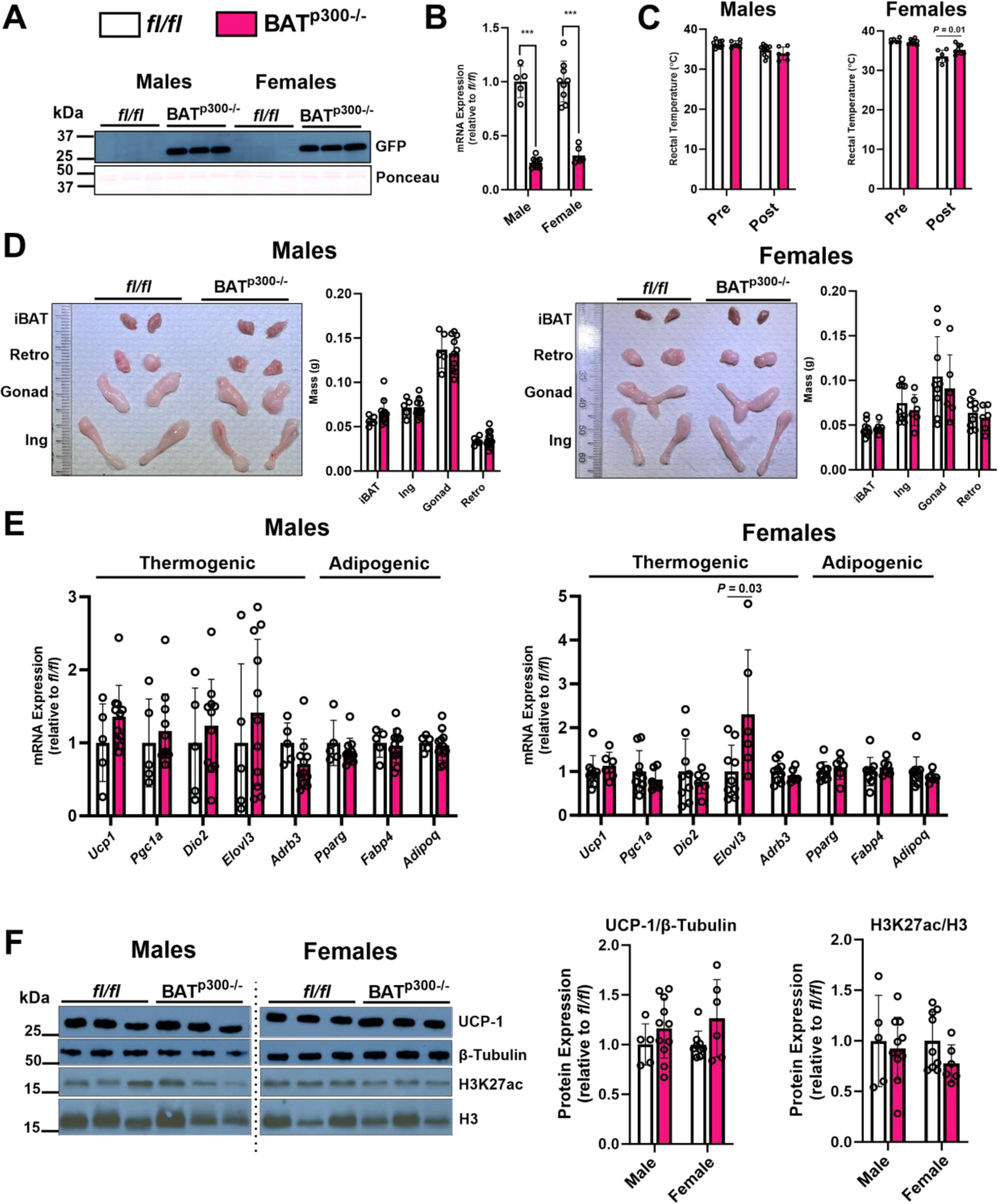
Loss of p300 does not impair BAT formation of function *in in vivo*. **A)** Positive expression of green fluorescent protein in male and female BAT^p300^–^/-^ only. Equal protein load was confirmed by Ponceau stain. **B)**. Knockdown of brown fat *ep300* exon 9 gene expression. **C)** Intrarectal temperature (°C) before (pre) and after (post) exposure to 4°C for 6 hrs. **D)** Tissue wet weights (g) and representative photographs of interscapular brown fat (iBAT), inguinal (Ing), gonadal (Gonad), and retroperitoneal (Retro) white adipose depots. **E)** Brown adipose tissue thermogenic and adipogenic marker gene expression. Transcripts were normalized to *Actb* and expressed relative to *fl/fl* controls. **F)** Interscapular brown adipose tissue protein expression. Values are Avg. ± S.E.M. ** *P* < 0.001. n = 5-10/group.

### Diet-Induced Obesity

While integral for adaptive thermogenesis, brown adipose tissue plays other roles in regulating peripheral metabolism via production and release of BATOkines, some of which are important regulating fatty acid oxidation and glucose metabolism,^17–20^ among other functions. Furthermore, BAT thermogenesis can be modulated by calorie surfeit in rodents (i.e. diet-induced thermogenesis).^1^ Given its role in co-ordinating transcriptional activation of gene programs, we next determined whether p300 HAT activity was obligatory for activating diet-induced metabolic programming of brown fat. To do so, mice were given a “Westernized” high-fat diet for 16 weeks. Weight-gain was comparable between *fl/fl* and p300^BAT-/-^ littermates throughout the 16-week HFD (**Fig. 2A**). Consistently, body composition (i.e. total lean/fat mass) were comparable between genotypes immediately prior to, after 8 and 16 weeks of diet-induced obesity (**Fig. 2B**), suggesting diet-induced thermogenesis remained intact in p300^BAT-/-^ mice. Furthermore, glucose tolerance was not different between control and knockout mice prior to and throughout the HFD (**Fig. 2C**), indicating that BAT-specific loss of p300 did not affect glucose handling independent of adiposity.

**Figure 2.**
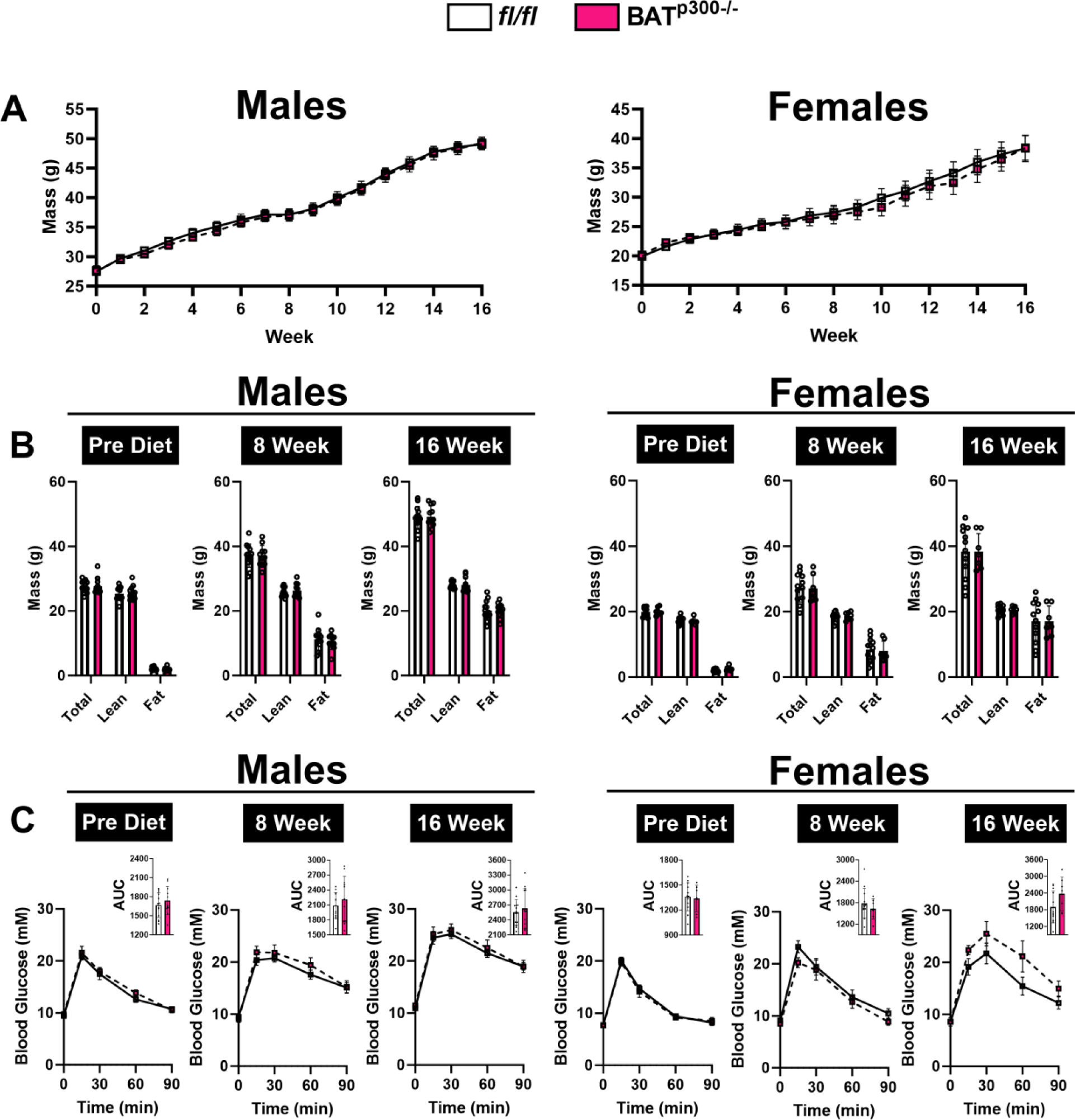
BAT-specific loss of p300 does not predispose mice to diet-induced obesity. **A)** Body mass (g) throughout the course of the 16 week “Westernized” high-fat diet. **B)** Body composition (total, lean, fat mass) immediately prior to (Pre-Diet) and throughout (8, 16 Week) the diet. **C)** Intraperitoneal glucose tolerance of mice throughout the dietary course. Inset images display area under the glucose curve (AUC). Values are Avg. ± S.E.M. n = 7-13/group.

### “Browning” of White Adipose Tissue by a Selective β_3_-Adrenergic Receptor Agonist

Signalling through the β_3_ isoform of the adrenergic receptor (AR) is critical for acutely activating BAT and for enhancing its thermogenic capacity.^1^ Furthermore, β_3_-AR agonism stimulates *de novo* biogenesis of beige adipocytes within subcutaneous fat (i.e. “browning” of white fat). While our previous experiments revealed p300 expression to be dispensable for classical brown adipose, we next sought to examine its requirement for the induction of beige adipocytes, specifically because they arise from a distinct cellular lineage than that of brown adipocytes. To test the requirement of p300 in the “browning” of white fat, mice were given daily intraperitoneal injections of the highly-selective β_3_-AR agonist CL-316,243 or saline vehicle for 7 days. As expected, BAT^p300^–^/-^ showed a significant depletion of inguinal adipose *ep300* exon 9 mRNA, although this genotype effect was less pronounced in females (**Fig. 3A**). While knockdown was lower in magnitude to that in BAT (**Fig. 1B**), inguinal WAT contains considerably more non-adipocytes than classical interscapular brown fat.^41^ Furthermore, GFP expression was markedly induced in CL-treated BAT^p300^–^/-^ mice (**Fig. 3B**), indicating successful recombination within newly formed beige adipocytes specifically. Despite loss of p300, expression of thermogenic marker genes were greatly induced by CL treatment, irrespective of mouse genotype (**Fig. 3A**). As expected, UCP-1 and mitochondrial respiratory chain complex protein expression were completely absent or nearly undetectable in saline treated ING depots, but markedly induced by CL administration (**Fig. 3B**). Given this, we chose to compare their expression in CL-treated mice only. Consistent with our gene expression findings, UCP-1 protein content and respiratory chain components were induced equivalently between control and knockout mice, suggesting *de novo* beige adipogenesis remained intact despite loss of p300 from thermogenic fats. Interestingly, complex IV expression was lower in CL-treated BAT^p300^–^/-^ females (**Fig. 3B**).

**Figure 3.**
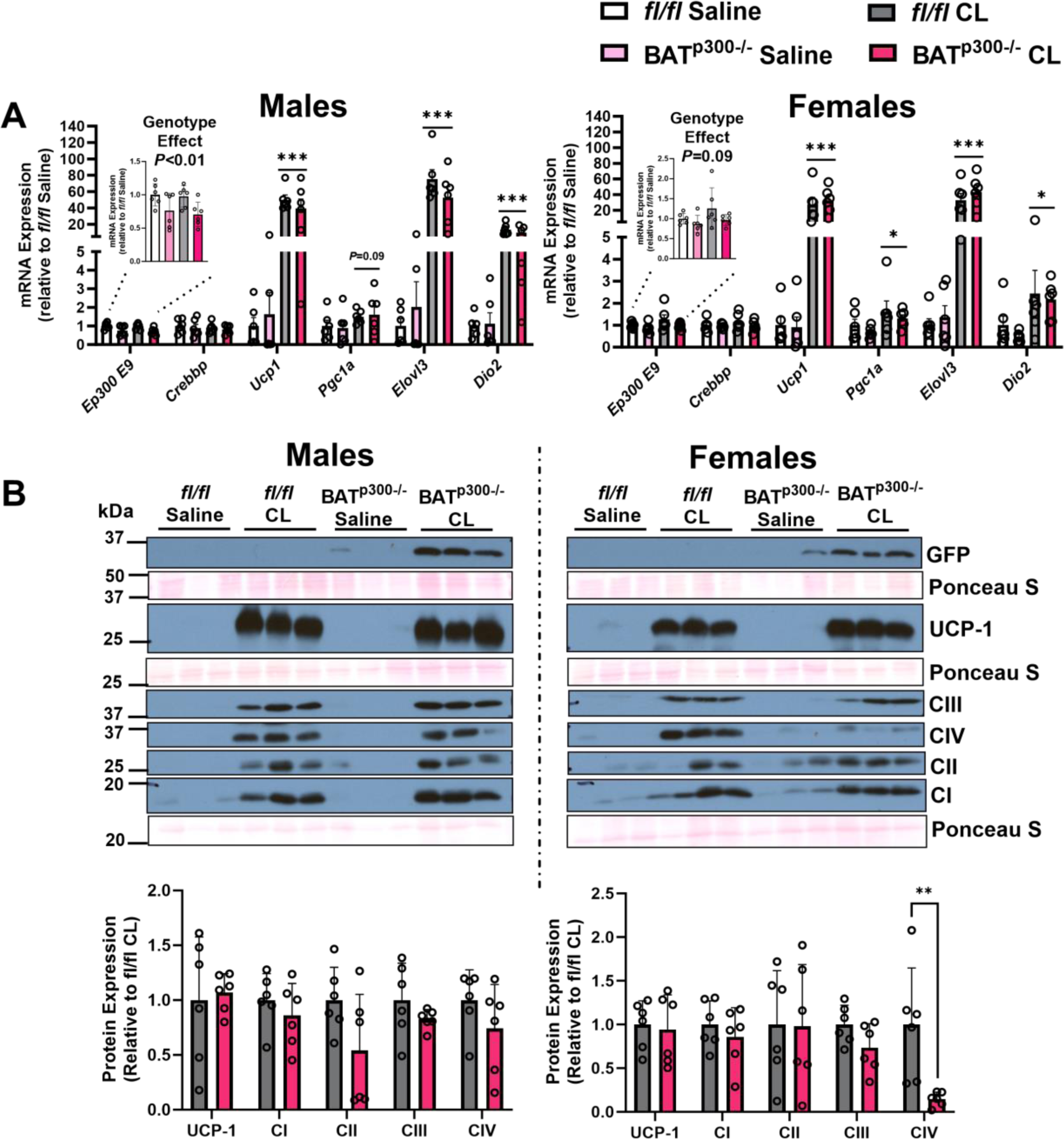
“Browning” of subcutaneous white fat is unaffected by p300 loss. **A)** Expression of thermogenic marker genes of the inguinal white adipose depot. Inset displays depletion of ep300 exon 9 (E9) expression in BAT^p300^–^/-^ animals. Main effect of CL treatment: * *P* < 0.05, *** *P* < 0.001 **B).** Inguinal white adipose protein expression of saline and CL treated mice. Proteins were normalized to their corresponding Ponceau S stains below. ** *P* < 0.05. Values are Avg. ± S.E.M. n = 6/group.

## Discussion

Previous studies of histone deacetylases have suggested the transcriptionally activating H3K27ac mark is critical for functional homeostasis and adaptability of both brown and beige adipose tissues.^22–28^ Here, we tested the hypothesis that p300, an epigenetic writer responsible for adding acetyl groups onto H3K27, would be required for brown and beige adipose programming *in vivo*. Using *Ucp1-Cre* to selectively remove the *ep300* gene from thermogenic fats (leaving white adipose intact), we show that BAT^p300^–^/-^ mice retained their ability to form and maintain classical brown fat mass and functionality. As such, BAT^p300^–^/-^ mice were not susceptible to diet-induced obesity or dysregulation of whole-body glucose handling. Furthermore, the absence of p300 did not prevent *de novo* biogenesis of beige adipocytes within the inguinal depot (i.e. “browning” of white fat). Together, our findings indicate that the activity of p300 specifically is dispensable for thermogenic gene programming of murine brown and beige fats.

Unlike other adipose-selective Cre systems (e.g. *Adipoq*-Cre, *Fabp4*-Cre),^42^ we chose *Ucp1*-Cre for our studies because it would target recombination within brown and beige adipocytes specifically.^43^ Furthermore, it has been used to create viable BAT-specific knockouts of several histone modifying enzymes.^23,44–46^ Because *Ucp1* expression is detected abundantly in late fetal development (i.e. E19) and maintained into adulthood,^47^ we expected p300 knockout to markedly impair the developmental expansion of brown fat, resulting in a persistent impairment of thermogenic function into adulthood of knockout mice; such a phenotypic trajectory has been observed when knocking out other epigenetic regulators important for transcriptional activation with *Ucp1-*Cre.^23,44,45^ Given its role in chromatin relaxation, we expected loss of p300 to markedly impair activation of thermogenic gene programming in BAT. Despite successful knockdown, BAT^p300^–^/-^ showed no impairment in whole-body energy metabolism, acute cold-tolerance, or expression of thermogenic marker genes and proteins (e.g. UCP-1). Interestingly, female knockouts were better able to defend their body temperature during thermal challenge, consistent with their higher levels of brown fat *Elovl3* expression. *Elovl3* encodes for elongation of very long chain fatty acid family member 3, a cold-responsive C20-C24 fatty acid elongase.^48^ Given that brown fat primarily relies on fatty acid oxidation to fuel energy needs,^1^ an enhanced ability to synthesize fatty acid substates would better support thermogenesis during cold-challenge. Currently, it is unclear how loss of p300 enhances *Elovl3* expression in females only, although such a mechanism could result from derepression of *Elovl3* transcriptional regulators dependent on p300 HAT activity.

Despite lack of a major H3K27 acetyltransferase, BAT^p300^–^/-^ fully retained their H3K27ac levels, as measured by Western blot. This ability is likely due to functional compensation from p300’s homologue, CBP. Our results are consistent with a previous report showing that single gene knockout of either p300 or CBP from mouse embryonic fibroblasts is inconsequential to global H3K27ac content.^35^ Studies of p300 specifically in skeletal muscle^49^ and across all adipose tissue types^50^ have found its absence inconsequential for tissue homeostasis and function. Thus, the presence of CBP in BAT^p300^–^/-^ mice likely provides sufficient functional overlap to maintain gene programming, preserving brown fat development and metabolism under standard rodent housing conditions.

Brown adipose metabolism can be altered by various conditions including prolonged high-fat feeding, which increases UCP-1 expression.^1,40^ Furthermore, numerous rodent studies have shown that impairing brown adipose tissue metabolism leads to obesity and metabolic dysfunction in room temperature housed mice. While our initial experiments revealed no major defects in BAT^p300^–^/-^ mice, we could not rule out a contextual requirement of p300 HAT activity, particularly during states known to remodel brown fat metabolism that would require transcriptional activation of adaptive gene programs. As such, mice were fed an obesogenic diet to see if a phenotype would emerge during prolonged calorie surfeit. Consistent with our initial metabolic characterizations, BAT^p300^–^/-^ mice were not susceptible to diet-induced obesity or dysregulated glucose handling as hypothesized, suggesting diet-induced recruitment of UCP-1 and energy balance were unaffected by p300 knockdown. Furthermore, our results suggest that BATokine programming, specifically those important for peripheral glucose handling,^17^ also remained intact in knockout mice. Consistent with our findings, pan-adipose deletion of either p300 or CBP using *Adipoq*-Cre has no effect on body composition, glucose or insulin tolerance.^50^ However, knockout of both HATs with this Cre system causes generalized lipodystrophy, hyperglycemia, hyperlipidemia, and hepatomegaly.^50^ Thus, in the absence of p300, CBP affords enough functional overlap to maintain brown adipose H3K27ac distribution to support BAT adaptability in the context of diet-induced obesity.

Unlike classical brown fat whose development arises prenatally from *Myf5*+ precursor cells of the dermomyotome, beige adipocytes have a distinctive cellular origin and are induced postnatally by cues like cold exposure, hormones, or even during diseases like cancer cachexia.^6^ While p300 is dispensable in the formation and function of tissue arising from a *Myf5*+ origin like skeletal muscle^49^ and BAT (show here), it was unclear if this was also the case for *de novo* biogenesis of beige adipocytes *in vivo*. We chose to examine this pharmacologically by treating animals with the highly selective β_3_-AR agonist CL-316,243 and examining thermogenic markers within inguinal subcutaneous fat, which has a high propensity for browning.^6^ We found that expression of thermogenic genes, UCP-1 protein, and most mitochondrial respiratory chain components were induced comparably by CL treatment in BAT^p300^–^/-^ relative to littermate controls. Our results suggests p300 is not obligatory for beige adipogenesis, and is supported by previous work that cold-induced thermogenic gene expression profiles of inguinal fat are unaltered by pan-adipose p300 deletion.^50^ As is the case for brown fat, it is likely that functional compensation by CBP is sufficient to maintain beige adipogenesis *in vivo* despite p300 knockout.

Brown and beige fats play an integral role in thermoregulation and systemic metabolism in mice. Although a wealth of rodent studies showing that enhanced BAT and beige activity protects against obesity, adult humans have relatively little active brown fat.^13^ Despite this, human BAT activity is inversely correlated with body mass index and fasting blood glucose.^10–12,16^ As such, enhancing the functional capacity of brown/beige fat could help mitigate metabolic dysfunction. However, our understanding of molecular factors important for establishing, maintaining, and controlling the adaptability of thermogenic fats are incompletely understood. Mechanisms governing chromatin state, therefore gene programming, are integral in this respect. This is the first report examining the requirement of the H3K27 acetyltransferase p300 within thermogenic adipose tissues specifically. We show here that p300 HAT activity is completely dispensable for the formation and activation of brown and beige adipose tissues *in vivo*, likely due to the presence of its functional homologue CBP.

## Supplementary Figures/Tables

**Figure S1.**
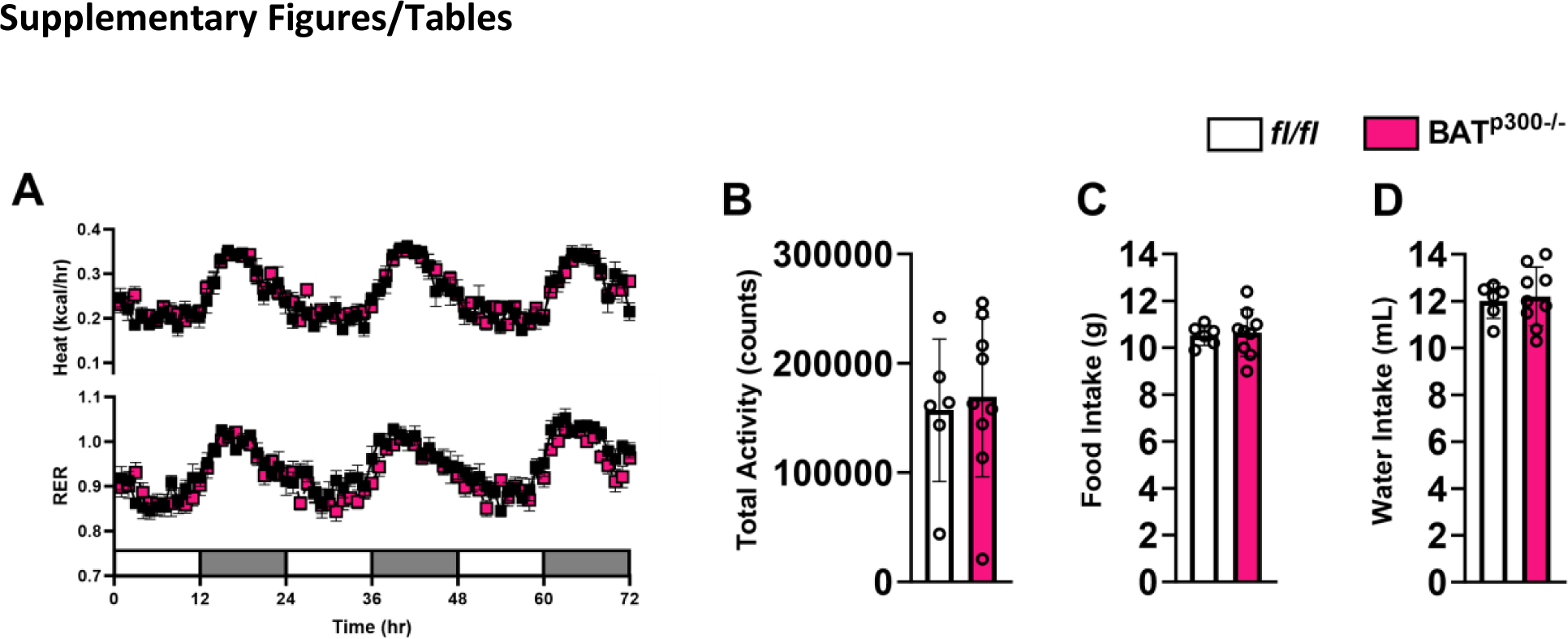
Whole-body metabolic phenotyping of female using the PhenoMaster metabolic cage system. **A)** Total energy expenditure and respiratory exchange ratio (RER) measured over 3 consecutive days. 12/12-hr light/dark cycle is depicted by unshaded and shaded regions of the x-axis, respectively. **B)** Cumulative cage ambulation (counts) measured by infrared beam breaks. **C)** Cumulative food (g) intake. **D)** Cumulative water (mL) intake. Values are Avg. ± S.E.M. *fl/fl* n = 6, BAT^p300^–^/-^ n = 9.

**Table S1.**
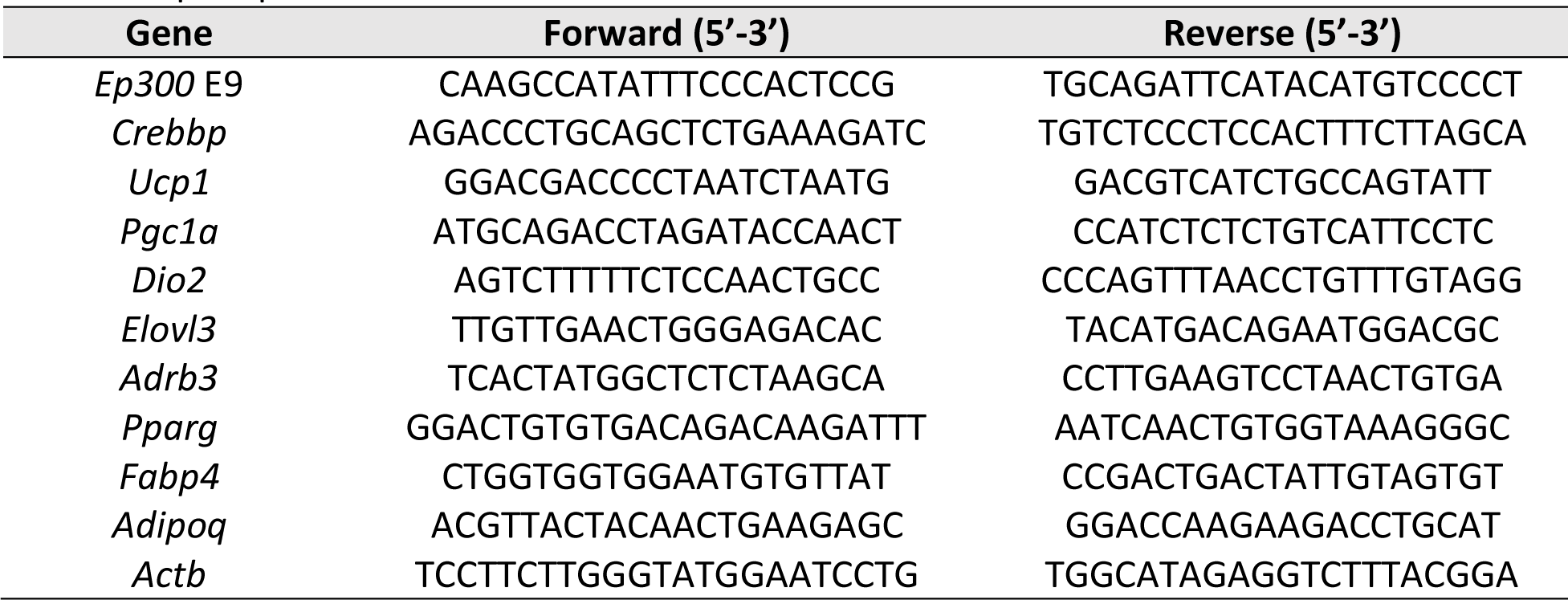
qPCR primer sets.

